# Towards a Dataset for State of the Art Protein Toxin Classification

**DOI:** 10.1101/2024.04.14.589430

**Authors:** Chance A. Challacombe, Nikhil S. Haas

**Affiliations:** Machine Learning Scientist at BioLM.ai; Cofounder at BioLM.ai

**Keywords:** machine learning, toxin classification, fine-tuning, computational biology, computational toxicology, biosecurity, peptide therapeutics, protein engineering, synthetic biology, protein language models, fine-tuning

## Abstract

*In-silico* toxin classification assists in industry and academic endeavors and is critical for biosecurity. For instance, proteins and peptides hold promise as therapeutics for a myriad of conditions, and screening these biomolecules for toxicity is a necessary component of synthesis. Additionally, with the expanding scope of biological design tools, improved toxin classification is essential for mitigating dual-use risks. Here, a general toxin classifier that is capable of addressing these demands is developed. Applications for *in-silico* toxin classification are discussed, conventional and contemporary methods are reviewed, and criteria defining current needs for general toxin classification are introduced. As contemporary methods and their datasets only partially satisfy these criteria, a comprehensive approach to toxin classification is proposed that consists of training and validating a single sequence classifier, BioLMTox, on an improved dataset that unifies current datasets to align with the criteria. The resulting benchmark dataset eliminates ambiguously labeled sequences and allows for direct comparison against nine previous methods. Using this comprehensive dataset, a simple fine-tuning approach with ESM-2 was employed to train BioLMTox, resulting in accuracy and recall validation metrics of 0.964 and 0.984, respectively. This LLM-based model does not use traditional alignment methods and is capable of identifying toxins of various sequence lengths from multiple domains of life in sub-second time frames.

## Introduction

Toxin classification is an important target for machine learning in computational biology with important applications in both industry and research settings, particularly for genetic engineering, protein and drug design, biosecurity and toxicology. For instance, genetic engineering often involves expressing an introduced or modified protein; consequently screening these proteins for toxic effects (towards the target organism or wider populations, e.g. humans) is important. Also, in the case of genetically engineered (GE) crops, this process is required before commercialization (1). *In-silico* toxin classification is an essential piece of this screening process. Proteins and peptides are increasingly prevalent options for therapeutic development (2, 3) and have applications in modulating protein protein interactions (PPI), pain, cancer and diabetes. Currently, more than 80 peptide drugs approved worldwide with more than 170 are in development, and these numbers are expected to increase (2). As with any new medical applications, it is critical that these proteins and peptides are tested for potentially unexpected or dangerous effects, and in order to gain FDA approval, new therapeutics must possess an acceptable safety profile (4). While *insilico* toxin classification is not currently a direct replacement for laboratory testing, it is much faster and less resource intensive, allowing companies and researchers to iterate their designs before employing more expensive validation methods.

Recent enhancements in machine learning have created opportunities for improvement on many computational biology tasks, including toxin classification. Simultaneously, these enhancements have amplified both the capabilities and dual-use risks (potential risks associated with technology that can be used for both beneficial or malicious purposes) of biological design tools (BDTs) (5). BDT’s are increasingly able to assist in protein engineering and design (6–9). Consequently, whether inadvertently or with malicious intent, BDT’s may create dangerous biomolecules. In a 2022 wake up call and report, Urbina et al. (10) detailed an example of possible, dangerous, *de novo* design of toxins using machine learning BDTs, demonstrating that limiting and reducing access to datasets that contain dangerous biomolecules may not entirely erase these risks. Beyond just generating potentially toxic biomolecules, D’Alessandro et al. (11) suggest that eventually, if not currently, BDTs will be capable of creating threats that escape standard biosecurity and laboratory security detection.

Verkuil et al. (12) have shown that language models can access a design space beyond that of natural proteins, designing novel sequences and structures that can be successfully expressed, a capacity that may open new opportunities in protein design. This raises critical challenges for synthesis screening as functional proteins with low sequence homology to known proteins become increasingly prevalent (13). Since many sequence alignment and similarity based detection methods are unlikely to identify novel toxins that are dissimilar in structure or sequence to known toxins, taxonomy and sequence based detection methods will likely no longer be adequate for many biosecurity tasks (5). As BDTs become increasingly capable of creating difficult to detect toxins, high performing general toxin classifiers become an immediate necessity.

Academically, toxin classification is an important step for computationally determining mechanisms of toxicity. As classifiers are more able to determine what a toxin is, the problem of determining mechanisms of toxicity and eventually toxin specificity (14) becomes more accessible. Toxin classification is also interesting from an evolutionary perspective. Toxin homologs are expressed in low to moderate levels in non-secretory tissues besides the venom gland of snakes and can lead to confounding evolutionary analyses (15, 16). Accurate classification of these homologs with computational methods may increase the speed and confidence of these analyses. In fact, venom protein homologs appear to be broadly expressed throughout many tissues of both venomous and non-venomous snakes, suggesting that the evolution of venom toxins descends from genes with normal physiological roles (16). Researchers could rapidly test theories about which mutations initially transformed these physiological proteins into toxin candidates by screening mutated sequences with a toxin classifier.

Additionally, several proteins including lipocalin, phospholipase A2 (type IIE) and vitelline membrane outer layer proteins, have been hypothesized to be new toxins from rear fanged snake species, but have not been functionally characterized in venoms (15). Toxin classification with a computational model is a fast and inexpensive way to evaluate hypotheses, potentially motivating or even deprioritizing efforts for experimental validation.

While screening novel or modified proteins and peptides for potentially dangerous and toxic effects is an important step in verification of therapeutic design, venoms and toxins themselves have been recognized as an incredible source of bioactive peptides, incurring interest as potential bases for treatment options (17–19). The division between helpful and harmful is subtle and an expanding area of interest that may be enhanced with computational toxin classifiers.

### Conventional Methods

One of the most straightforward approaches to toxin classification has been sequence similarity or alignment methods (20) such as the Levenshtein distance or a tool like NCBI’s BLAST (21). While BLASTP (BLAST for proteins) has been used a baseline for toxin classification methods, due to the influx of new classifiers that outperform BLAST alone (22, 23), BLAST is no longer considered a state of the art (SOTA) method. Another similarity based alternative for protein function classification is Locality-Sensitive Hashing (LSH), which uses vector representations of sequences that are hashed repeatedly (24), and is generally a much faster approach than BLAST (25). However, as venoms and toxins frequently evolve from non-toxic proteins, sequence similarity based methods may struggle with paralogs (20, 26).

More recently, machine learning methods have been used as replacements for, or in conjunction with, alignment based methods. A naive feature extraction method for toxin classification is one-hot encoding, a heuristic that generates sparse sequence representations of features without capturing some relationships such as the similarity between polar residues. Another feature extraction method is bag of words (BoW); BoW is used to count the composition of a sequence either for individual tokens (for proteins these can be amino acids), combinations of tokens without overlap (n-grams) and combinations of tokens with overlap (k-mers) (24). This method, since it relies on counting, discards information about the order of amino acids in the sequence.

Historically, machine learning based toxin classifiers have used feature arrays of physio-chemical properties, evolutionary profiles, and/or molecular graphs. However, because a protein is defined by its underlying amino-acid sequence, biological information, is necessarily encoded in the sequence (24, 27) which has led to NLP based approaches for feature extraction from sequences to be considered.

### Contemporary Methods

The transformer architecture introduced by Vaswani et al. (28), is a staple of natural language processing (NLP) and the basis of large language model (LLM) technologies, and in particular has shown capabilities in extracting structural, binding site and additional biophysical information from protein sequences (12, 29). For these reasons, sequence language models and NLP methodologies can be used for toxin classification without the need for alignment or additional features.

Contemporary toxin classification methods that make use of large language models are UniDL4BioPep (30) and CSM-Toxin (31). UniDL4BioPep uses pre-trained protein language model embeddings as features into additional deep learning layers. CSM-Toxin is a fine-tuned iteration of the pre-trained ProteinBERT (32) language model. Additional contemporary methods for toxin classification use features including molecular graphs, physiochemical properties, and evolutionary information from PSSM and BLOSSUM substitution matrices (these require alignment). The architectures for these methods include convolutional neural networks (CNNs), gated recurrent units (GRUs) and graph neural networks (GNNs) (Table 1). These methods are TOXIFY (26), ToxVec (33), ToxDL (22), ToxinPred2 (23), ATSE (34), ToxIBTL and ToxIBTL Variational Information Bottleneck (ToxIBTL VIB) (35, 36). Contemporary Toxin classification models achieve competitive results, though sometimes they are high performing in a limited scope.

**Table 1.** Overview of contemporary toxin classifier methods. The architecture column specifies the architecture for machine learning components of these classifiers. The domains of life column specifies the taxonomic origin of sequences the models for each method used for training. The features column details the features and input for each method. The special evaluation column describes any special evaluations that were performed that pertain to general toxin classification or the proposed criteria

| Method | Architecture | Domains of Life | Sequence Length | Features | Special Evaluations for General Toxin Classification |
| --- | --- | --- | --- | --- | --- |
| TOXIFY<br>Cole and Brewer (26) | Gated Recurrent Units (GRUs) | Eumetazoan venoms from Swissprot | $\leq 500$ AA's | Atchley factor matrices | Non toxic homologs for Robber Fly toxin |
| ToxVec<br>Ahmadi et al. (33) | Fasttext model | Eumetazoan venoms from Swissprot | $\leq 500$ AA's | ProtVec embeddings of non overlapping 3-mer combinations | |
| ToxDL Pan et al. (22) | CNN with max pooling operations | Animal toxins from Animal Toxin Annotation Project | Contains sequences longer than 1002 AA's | One-hot encoded protein sequences and domain2vec embedding vectors | Toxic domain identification with integrated gradients and validation on bacterial sequences |
| ATSE Wei et al. (34) | Attention module on GNNs and CNN_BiLSTM |  | 10-50 AA's from SwissProt, ConoServer and ArachnoServer | Molecular graphs obtained by using RDKit and evolutionary information by PSSMs obtained by PSI-BLAST |  |
| ToxinPred2<br>Sharma et al. (23) | Ensemble ML Models, BLAST and a Motif based approach using MERCI | Multiple | $\geq 35$ AA's | Amino Acid Counts(AAC), Pfeature features, and PSSM profiles. These features are ranked and then reduced for each dataset | |
| ToxIBTL Wei et al. (35) | Hybrid CNN-BiGRU | Both ToxDL and ATSE datasets | Both ToxDL and ATSE datasets(longer proteins and short peptides) | BLOSUM62 evolutionary profiles and FEGS model generated graphical, statistical and physiochemical features | Analysis of peptide length biases for ToxIBTL and other SOTA methods |
| ToxIBTL VIB Zhao et al. (36) | Variation of ToxIBTL with a variational information bottleneck(VIB) | Same as ToxIBTL | Same as ToxIBTL | Same as ToxIBTL | Analysis of peptide length biases |
| UniDL4BioPep<br>Du et al. (30) | CNN | Same as ATSE | Same as ATSE | ESM-2 embeddings |  |
| CSM-Toxin<br>Morozov et al. (31) | Fine-tuned BERT LLM | | $\leq 5000$ from SwissProt | Base Pre-trained ProteinBERT model | |

### SOTA Criteria

Reviewing these contemporary methods, we identified five heuristic criteria for evaluating toxin classifiers as SOTA *general* toxin classifiers. These criteria encompass both dataset and architecture categories, and are discussed in relation to the contemporary methods below and in Table 1.

### Dataset Criteria

#### 1. Domains of Life

Some contemporary models are only trained on datasets of specific taxonomic origin. For example, the training datasets for ToxDL and TOXIFY include only animal venom sequences. Furthermore, there are more toxic proteins for certain taxonomies both within databases and amongst genomes that may cause taxonomy biases. TOXIFY addresses this by evaluating toxic and non toxic sequences from *Machimus arthriticus* (26).

#### 2. Sequence Length

Several contemporary models are only trained on datasets that contain sequences of a specific length. For certain methods (ATSE, UniDL4BioPep), high performance on peptides of shorter length was the target goal and generalizing to greater sequence length was not a criterion. However, generalizing to different lengths is an important capability. Length is also a consideration for some deep learning models. ATSE for example only uses input of size 50. Zero padding and truncation alleviates this limitation, however performance of this approach is not guaranteed. Furthermore, not all model evaluations include performance metrics on sequence length. ToxIBTL is one method that is evaluated on peptides in different length ranges, testing the model’s sequence length bias (35).

#### 3. Sequence Similarity and Homologs

A toxin classifier’s performance on proteins whose sequences are dissimilar to those in training is important. One potential strength of machine learning classifiers is that they can learn high level features for toxins that can be applied to proteins with dissimilar sequences. This is one of the reasons they have been employed either in conjunction with or as replacements for pure sequence similarity based methods like BLAST. Many of the contemporary methods discussed above use some variation of CD-Hit (37) to filter out proteins with similar sequences, to remove sequence similarity evaluation bias and to ensure their models can accurately predict on dissimilar sequences. This filtering method is often employed before the training and validation splits and can have the unfortunate effect of removing homologs between the prediction classes as well.

Toxin classifiers should be able to accurately distinguish between toxin and not-toxin homologs. If similar sequences are filtered out of training and validation datasets together, evaluation of the model on these homologs is no longer possible. Few methods perform this breakdown evaluation. TOXIFY is one method that does include an evaluation against not-toxin homologs. ToxDL uses CD-Hit to remove similar sequences to the validation set from the training set *after* splitting the data. This or a similar approach might allow evaluation of the model on different class homologs, however a more direct approach is to explicitly include selected homologs and dissimilar sequences in validation or to create separate datasets to allow these evaluations.

### Architecture Criteria

#### 4. Parameter and Feature Selection

One of the main reasons why sequence comparison methods like BLAST have become less favored is because thresholds and parameters must be selected in order to achieve a high performance. For this reason, the incorporation of feature selection and threshold learning into model architecture is desirable. In this regard, methods like ATSE, which make use of completely automated training and prediction without the need for a user to modify thresholds or manually select features, are very powerful. These methods can also be updated with new data and trained without needing to repeat feature selection.

#### 5. Computational Efficiency and Overhead

Transfer-learning and fine-tuning using a pre-trained model can be more computationally efficient than other methods that require training a model from scratch, as associations and features have already been learned. Contemporary methods that make use of these methods include ToxIBTL, UniDL4BioPep, ToxVec and CSM-Toxin.

Many methods also require feature generation that may include BLAST results, molecular graphs, physio-chemical properties, sequence statistical properties and evolutionary profiles. Feature generation, especially alignment based features, can increase inference and training time and contribute to model overhead. ATSE is noted to have time-consuming feature generation with PSI-BLAST (35).

Not all methods evaluate their model’s computational efficiency. Further complicating comparisons are hardware differences and model unavailability. Compute costs may also prohibit extensive comparison when there are many models to compare.

### BioLMTox

We propose BioLMTox, a single sequence general toxin classifier that demonstrates the performance capabilities of LLM fine-tuning for toxin classification with only minor prediction layer changes. BioLMTox was trained and validated on a unified dataset curated from a union of sequences from UniProt, UniRef and contemporary methods’ datasets. To the best of our knowledge, BioLMTox is the only toxin classifier that fine-tunes a large protein language model on a large representative dataset with explicit coverage over taxonomies from multiple domains of life and a wide range of sequence lengths Fig. 1, and is directly compared to contemporary methods using the validation datasets presented in those studies. Furthermore, during BioLMTox dataset development, we evaluated and removed ambiguously labeled sequences from the component datasets for a comprehensive comparison.

**Fig. 1.**
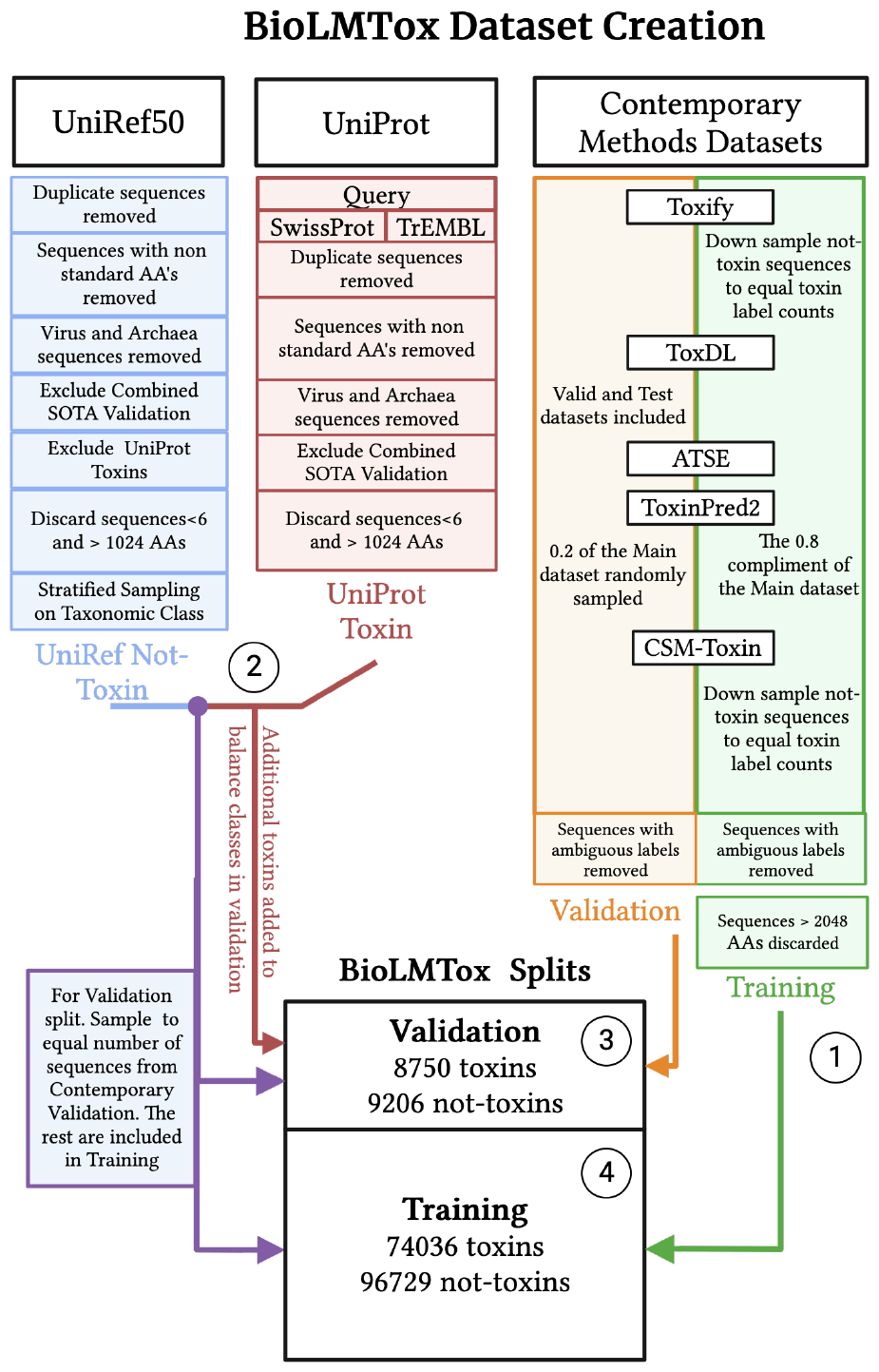
A diagram showing the creation and curation flow for the BioLMTox datasets. The four circled numbers in the diagram correspond to the BioLMTox dataset construction steps detailed in Sections A.1–A.4. Created with BioRender.com

## Methods

### Datasets

Motivated by the domain of life and sequence length SOTA data criteria, and in order to compare BioLMTox with the existing contemporary methods, we developed and curated a unified toxin classification dataset. Constructed to be a manageable size for fine-tuning while still maintaining ample validation data we reduce the amount of not-toxin data points (orders of magnitude more than the number of toxin sequences) with a variety of techniques and use selected data from eukaryotic and bacterial domains of life. Specific considerations were used to reduce bias and improve coverage over these domains, which include examples of amino acid sequences over a large range of lengths and removing sequences with conflicting labels amongst the contemporary method’s datasets.

The BioLMTox dataset were created with sequences from the UniProt and UniRef50 databases (38) and included datasets from other contemporary toxin classification methods. The inherent, drastic imbalance between toxin and not-toxin protein counts motivated the use of UniRef50, which contains representative sequences from a clustered non-redundant form of UniProt. Several downsampling steps were employed to reduce datasets to a reasonable size for fine-tuning while maintaining data representation across protein primary structure and taxonomies (stratified sampling on taxonomy). Including datasets from contemporary methods allows for direct validation compared to their respective methods. These datasets also contain sequences with characteristics that are useful for evaluating a general toxin classifier: ATSE’s dataset consists of short peptides, CSM-Toxin’s dataset contains much longer protein sequences, and Toxin-Pred2’s dataset is drawn from multiple domains of life. Figure 1 outlines BioLMTox dataset construction and Appendix Section A describes the explicit dataset construction methodology. As shown in Fig. 2, the majority of sequences in the BioLMTox dataset is selected from UniProt and UniRef as compared to the data selected from contemporary methods. The final toxin and not-toxin counts for the BioLMTox dataset is illustrated in Fig. 3.

**Fig. 2.**
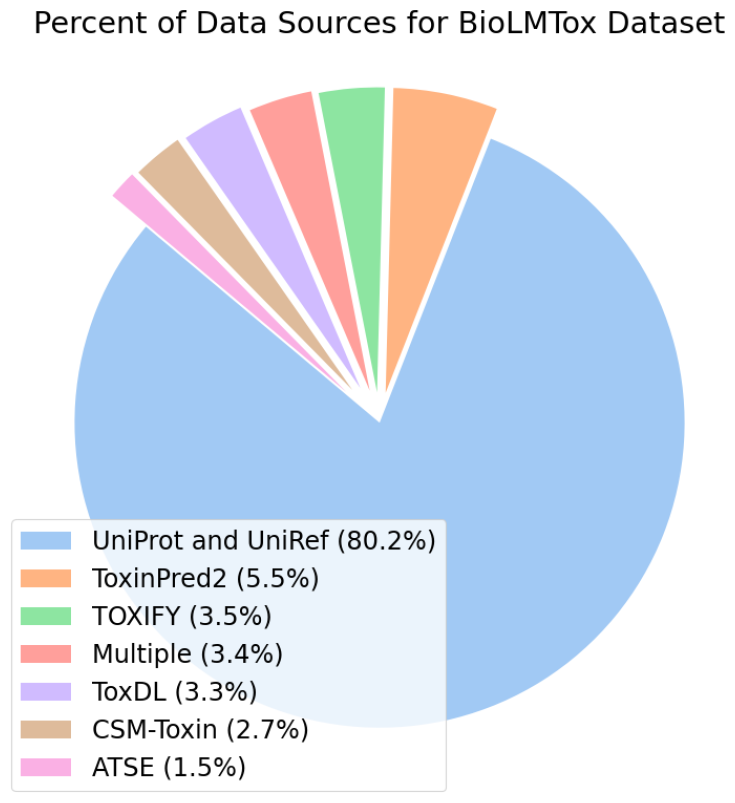
The dataset source frequencies for the entire BioLMTox dataset (train and validation). The Multiple label refers to overlapping data between datasets from two or more contemporary methods. UniRef and UniProt refers to the data from these sources selected during BioLMTox dataset creation

**Fig. 3.**
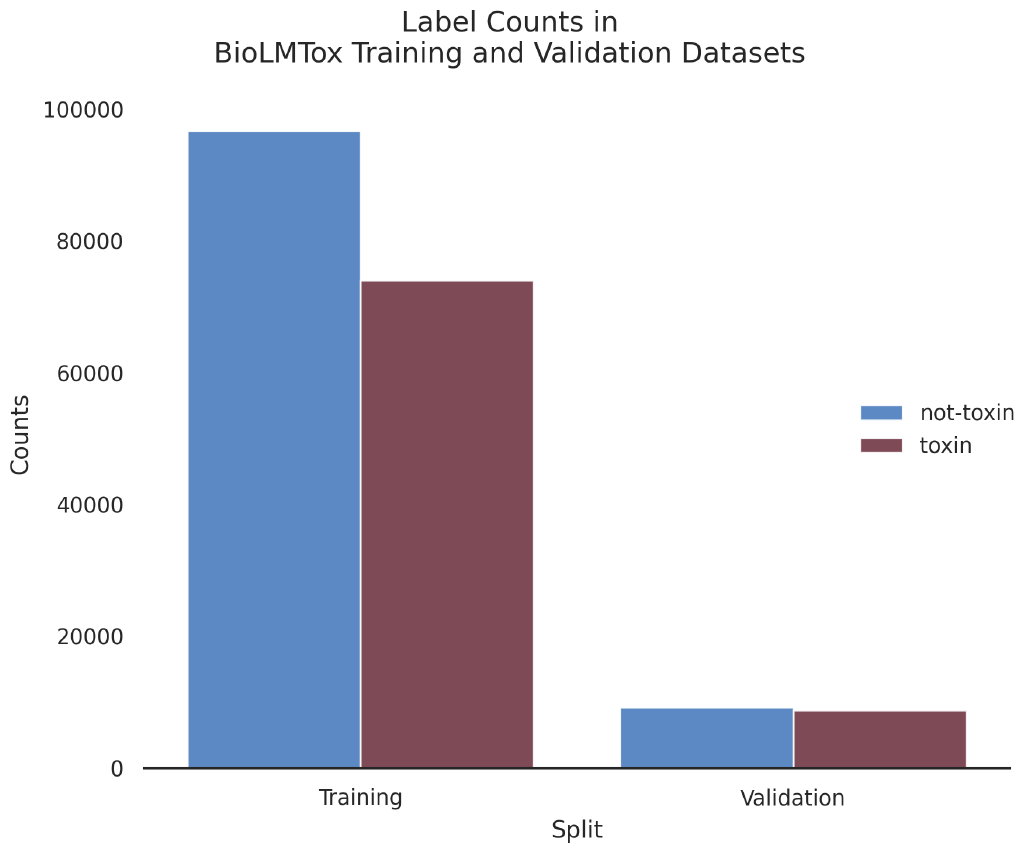
Label Counts for training (A.4) and validation (A.3) Datasets. The validation dataset contains 17956 total sequences with 9206 labeled as not-toxins and 8750 labeled as toxins. The final training dataset contains 170765 sequences with 96729 labeled as not-toxins and 74036 labeled as toxins

### Architecture

BioLMTox uses the esm2_t33_650M_UR50D pre-trained model with 650 million parameters and 33 layers. ESM-2 is a single sequence language model that was trained for masked language modeling (39). This means that it does not need any input in the form of evolutionary information, molecular graphs, physiochemical properties or any results from multiple sequence alignment (MSA). Instead, features are learned and extracted automatically from the sequence alone. The last layers of the BioLMTox architecture are a dense linear layer with dropout and tanh activation that takes the [cls] token hidden states of the pre-trained model as input (previous layers are not frozen, so pre-trained weights and layers are updated during training) and connects to another linear layer that outputs logits of dimension two (Fig. 4). Logits are unnormalized output vectors that determine the label prediction and can be used as input to the Softmax function to obtain the estimated probability for each predicted class. The Softmax function of the logit *x*_*i*_ with *n* labels and corresponding logits is defined with

**Fig. 4.**
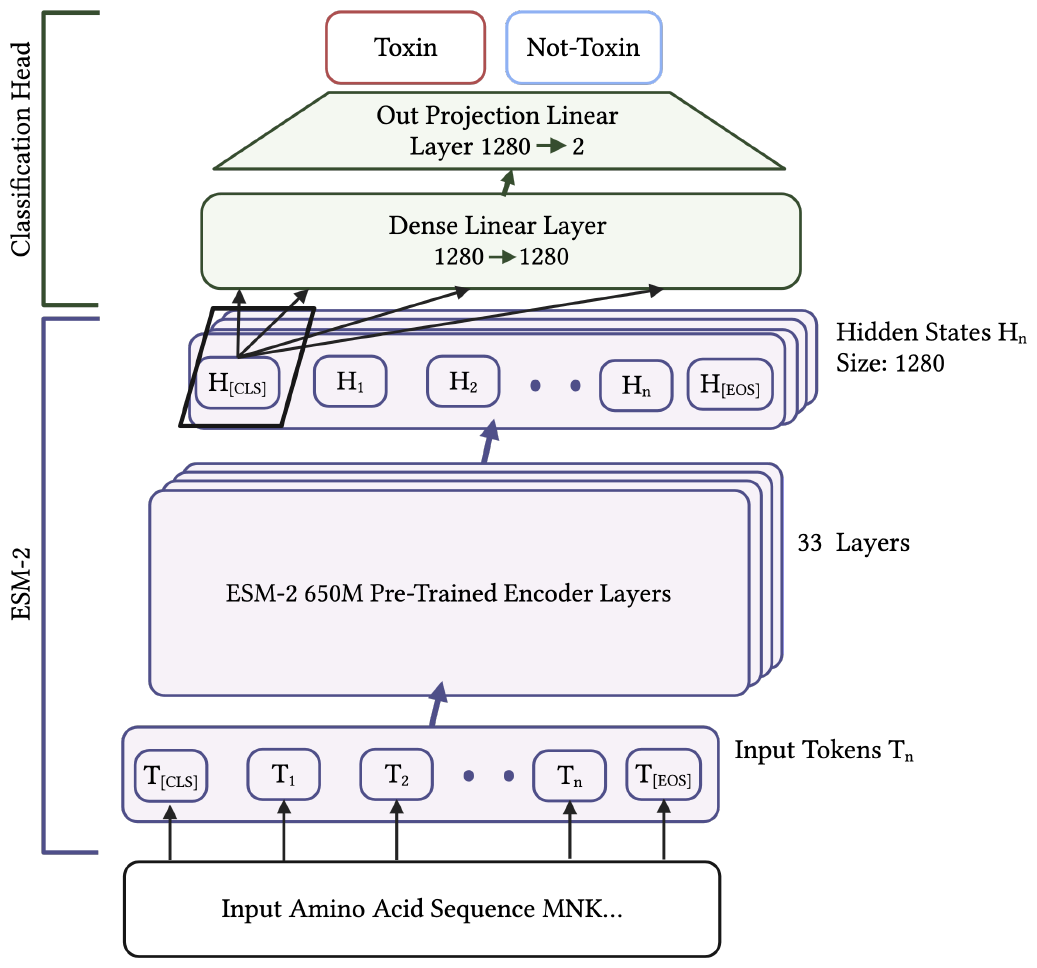
BioLMTox Fine-Tuning Architecture. Sequences are tokenized and a [CLS] (Classification) token is prepended to the residue tokens and an [EOS] (End of Sequence) token is appended to the residue tokens. These tokens are fed into the ESM-2 Encoder layers and last layers hidden states for these tokens is outputted. The [CLS] hidden states are then input into the added classification head to get a toxin or not-toxin prediction

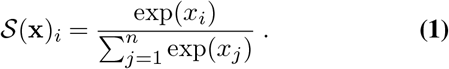

### Fitting and Tuning

The BioLMTox model was trained and instantiated using the HuggingFace Trainer (40) integrated with Ray Tune (41) for hyperparameter search using the ASHA algorithm (42).

During training, mixed precision, gradient accumulation and the ADAM 8 bit optimizer (43) were employed. The learning rate, gradient accumulation steps, warmup steps, and the optimizer’s *β*_1_, *β*_2_ and *ϵ* parameters were included in the hyperparameter search space. The loss function used during training was a weighted mean Cross-entropy that is widely used for classification tasks. The loss is given by

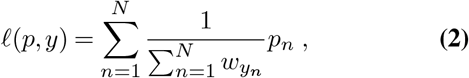

where *N* are the samples in the batch, *y* is the target (true) class, *w* is the class weight and *p*_*n*_ is the LogSoftMax of the logits *x*

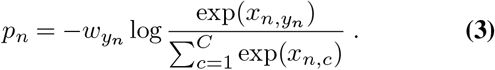

In the case where the class weights are all equal to one (2) becomes

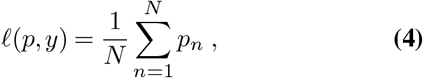

and (3) becomes

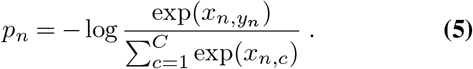

The model was fine-tuned using recall as the objective for two epochs with three different, randomly instantiated ASHA runs, which were then checkpointed and the best performing model for the recall objective was selected. This model, from that checkpoint, was further fine-tuned for an additional epoch with three more randomly instantiated ASHA schedulers, this time with accuracy as the objective. Figure 5a shows the training and evaluation loss curves during the last full ASHA runs for each instantiated scheduler.

**Fig. 5.**
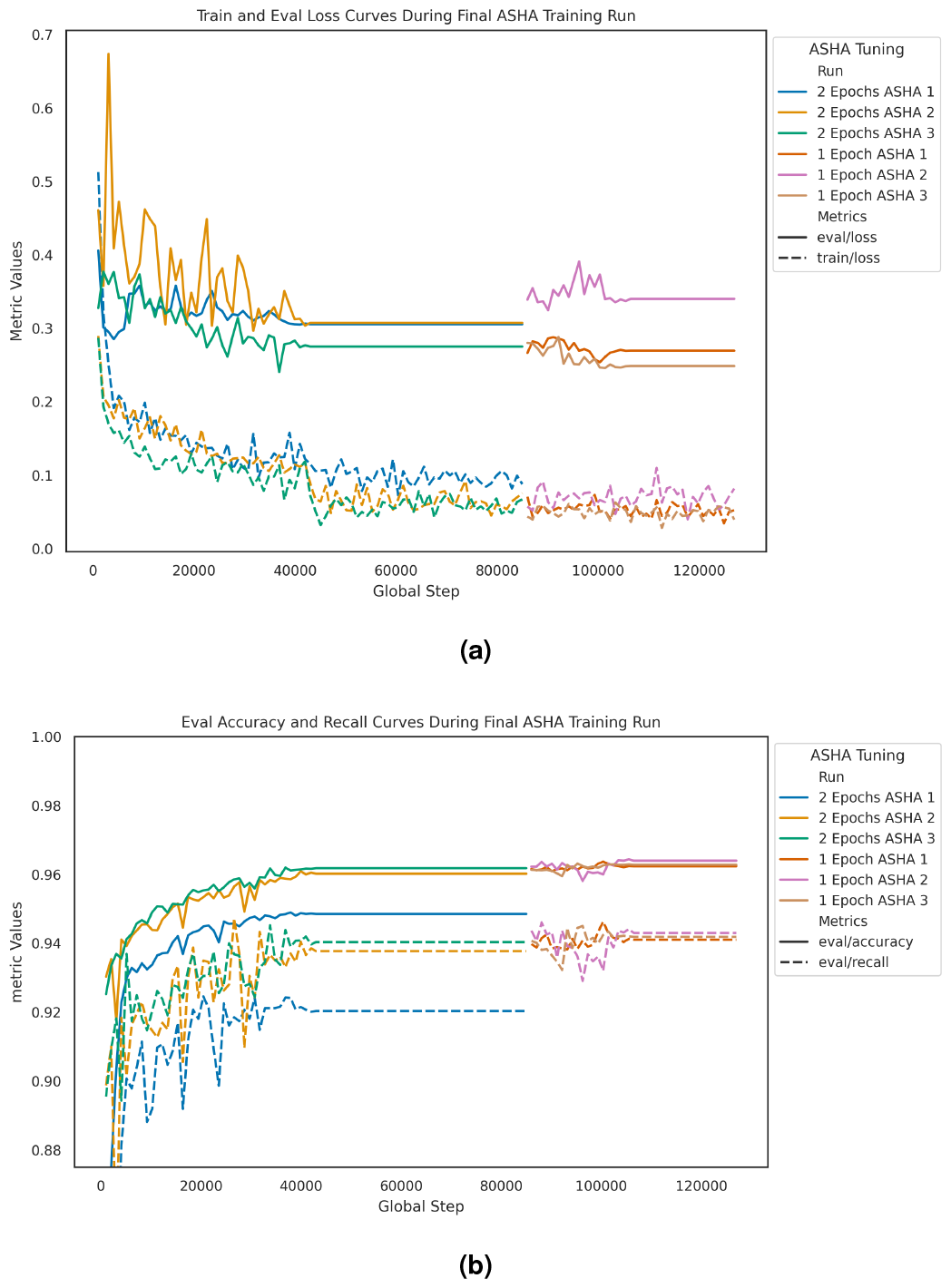
Metrics logged with WandB (44) during training of the final model (shown in different colors) for each instantiated ASHA scheduler. 5a) Train and Eval Loss curves during the ASHA runs. Train refers to the models metrics on the training dataset it is being fit to datasets A.3. Eval refers to the metrics on the validation dataset. The difference in line dash corresponds to eval or train for the loss curve. The eval loss curve, as expected, is above the training loss curve for the entire model fine-tuning since validation data is never used for training. The training and evaluation losses for the first iteration of ASHA schedulers continue to drop until the evaluation loss plateaus in the second epoch. Then, with the second iteration of the ASHA schedulers, variation is reintroduced into both loss curves. The evaluation loss eventually plateaus in this final epoch as well, with the training loss at 0.0524, 0.0816, and 0.0393 and the evaluation loss at 0.2694, 0.3401, and 0.2485 for each scheduler respectively. 5b) Eval Accuracy and Recall curves during the ASHA runs. While the loss for the 1 Epoch ASHA 2 model is greater than the other models, its accuracy of 0.964 is also greater than the other models at 0.9624 and 0.9628 (Fig. 5b).

### Model Selection

The model that yielded the best accuracy for the final epoch (the model trained with the second instantiated ASHA scheduler) was selected as the BioLMTox model (Fig. 5b).

## Results and Discussion

### Comparison with Contemporary Models

To directly compare BioLMTox’s performance against contemporary methods, BioLMTox predictions on the cleaned contemporary method datasets (Fig. 1. Appendix Section A) were evaluated. A side by side comparison of reported metrics for each contemporary method and BioLMTox metrics on the same dataset are recorded in Table 2 along with the percent difference in accuracy where applicable. As the Random Forest (RF) hybrid method was the best performing of ToxinPred2 hybrid methods on the method’s main dataset, this model was selected for comparison. Of the ToxIBTL variations, ToxIBTL VIB was selected for comparison. As detailed in Table 2, BioLMTox exceeds or is very near to matching accuracy on all methods except for ToxinPred2, where there is a -6.4 percent difference in accuracy. The ToxinPred2 dataset consists of all types of toxins available in the reviewed June 2 2021 UniProt release, and since BioLMTox was trained on a majority of bacterial and eukaryotic toxins (Fig.2), the model may not be generalizing to sequences of different origin. Additionally, the ToxinPred2 main dataset was noted to have conflicting labels with other datasets (A.1). This is possibly due to different definitions of toxin vs. not-toxin amongst datasets. Figure 6 illustrates that BioLMTox false negative (FN) on the ToxinPred2 dataset is higher than other methods, supporting this conclusion. There are 330 FN and 1314 true positives (TP) corresponding to a true positive rate (TPR) of 0.799, which is less than any of the breakdown TPRs for other contemporary methods.

**Table 2.** BioLMTox and reported metrics for the component contemporary datasets (A.1). The “Dataset” column indicates the origin dataset for both BioLMTox and the comparative methods (listed in the Methods column) predictions. Methods that are grouped between the same horizontal rules and have the same value in the “Dataset” column share the same dataset. ToxIBTL and ToxIBTL (VIB) were evaluated on two datasets that are shared with other methods and so are recorded twice (The ATSE dataset which consists of short peptides and the ToxDL dataset which contains longer proteins). The best metrics over all models for each dataset are boldfaced. The Acc %Diff column is the percent difference in accuracy between BioLMTox and the compared method.

| Contemporary |  | Contemporary Metrics |  |  |  | BioLMTox Metrics |  |  |  |  |
| --- | --- | --- | --- | --- | --- | --- | --- | --- | --- | --- |
| Dataset | Model | Accuracy | Recall | f1 | MCC | Accuracy | Recall | f1 | MCC | Acc %Diff |
| ATSE | UniDL4BioPep | 0.941 | 0.923 |  | 0.883 |  |  |  |  | −0.3 |
|  | ATSE | **0.952** | **0.965** |  | 0.903 |  |  |  |  | −1.5 |
|  | ToxIBTL | 0.9477 |  | 0.9428 | 0.9473 | 0.938 | 0.951 | 0.934 | 0.875 | −1.0 |
|  | ToxIBTL(VIB) | 0.949 |  | **0.9502** | **0.9483** |  |  |  |  | −1.2 |
| ToxDL | ToxDL |  |  | 0.809 | **0.793** |  |  |  |  |  |
|  | ToxIBTL(VIB) | 0.963 |  | 0.6775 | 0.6499 | 0.97 | 0.814 | **0.814** | 0.79 | 0.7 |
|  | ToxIBTL(VIB) | **0.9738** |  | 0.7528 | 0.7319 |  |  |  |  | −0.4 |
| TOXIFY | ToxVec | **0.95** |  | 0.89 |  |  |  |  |  | −1.7 |
|  | TOXIFY | 0.86 | 0.76 | 0.85 | 0.74 | 0.934 | **0.915** | **0.954** | **0.847** | **8.6** |
| ToxinPred2 | ToxinPred2 | **0.9554** | 0.9369 |  | 0.91 | 0.894 | 0.799 | 0.883 | 0.802 | −6.4 |
| CSM-Toxin | CSM-Toxin |  | 0.75 |  | 0.67 | 0.965 | **0.843** | 0.793 | **0.775** |  |

**Fig. 6.**
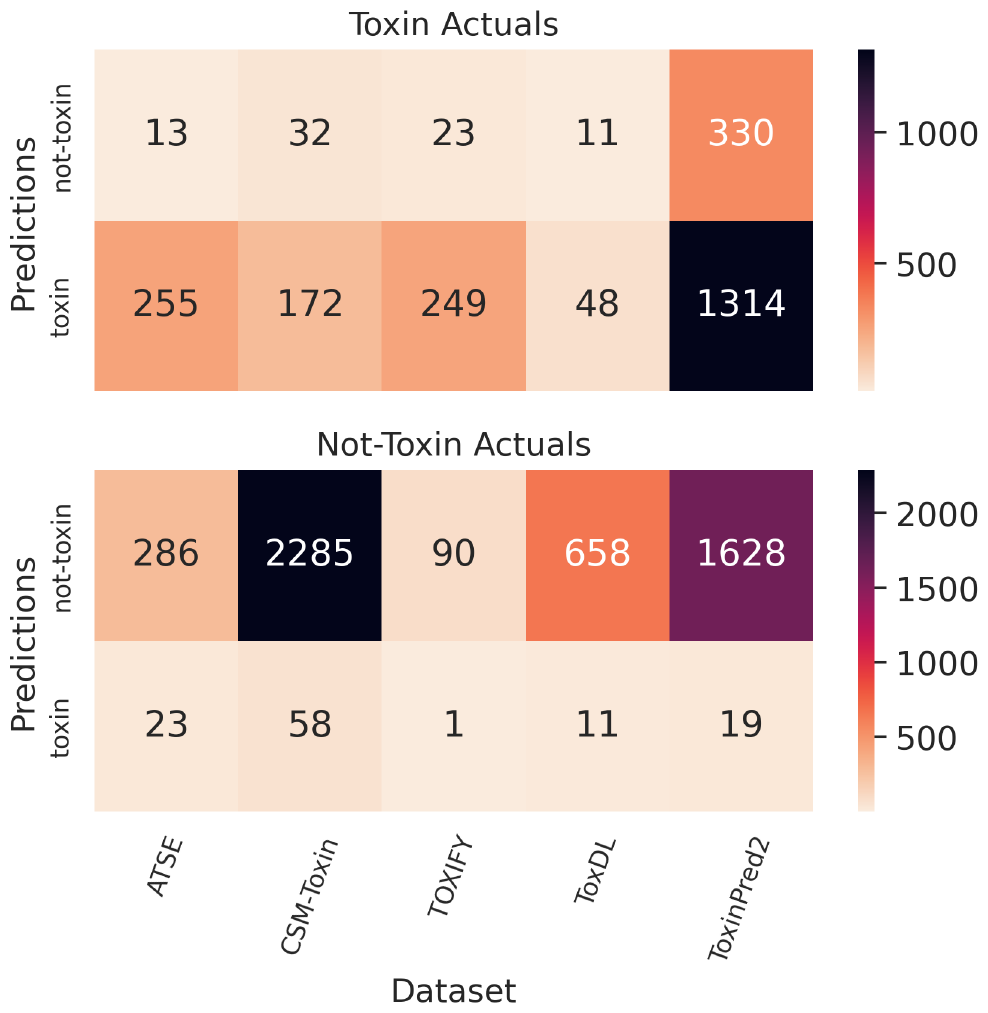
BioLMTox prediction results for each prediction class. Similar to a confusion matrix, true and false positives are shown in the top heatmap and true and false negatives are shown in the bottom heatmap. BioLMTox misclassifications are all relatively low compared to dataset size with the exception of ToxinPred2.

### BioLMTox Validation Dataset

Extending BioLMTox prediction evaluation from the contemporary methods datasets to the entire BioLMTox validation dataset, BioLMTox demonstrates strong classifier performance. The confusion matrix in Fig. 7 displays BioLMTox’s relatively small misclassification rate. While the FN (499) are greater than the FP (148), the ToxinPred2 dataset contributes more than half of the toxin FN misclassifications (Fig.6). Recorded in Table 3, BioLMTox achieved an accuracy of 0.964 and a Matthews Correlation Coefficient (MCC) of 0.929. Furthermore, the area under the receiver operating characteristic curve (auROC) and the area under the precision-recall curve (auPRC) are close to one at 0.986 and 0.989 respectively, indicating BioLMTox’s effectiveness as a classifier.

**Table 3.** BioLMTox Prediction Metrics for each prediction label (toxin and not-toxin) for the validation dataset (Datasets.A.3).

| Label | Accuracy | Recall | Specificity | Precision | f1 | MCC | auROC | auPRC |
| --- | --- | --- | --- | --- | --- | --- | --- | --- |
| not-toxin | 0.964 | 0.984 | 0.943 | 0.948 | 0.966 | 0.929 | 0.986 | 0.989 |
| toxin | 0.964 | 0.943 | 0.943 | 0.982 | 0.962 | 0.929 | 0.986 | 0.989 |

**Fig. 7.**
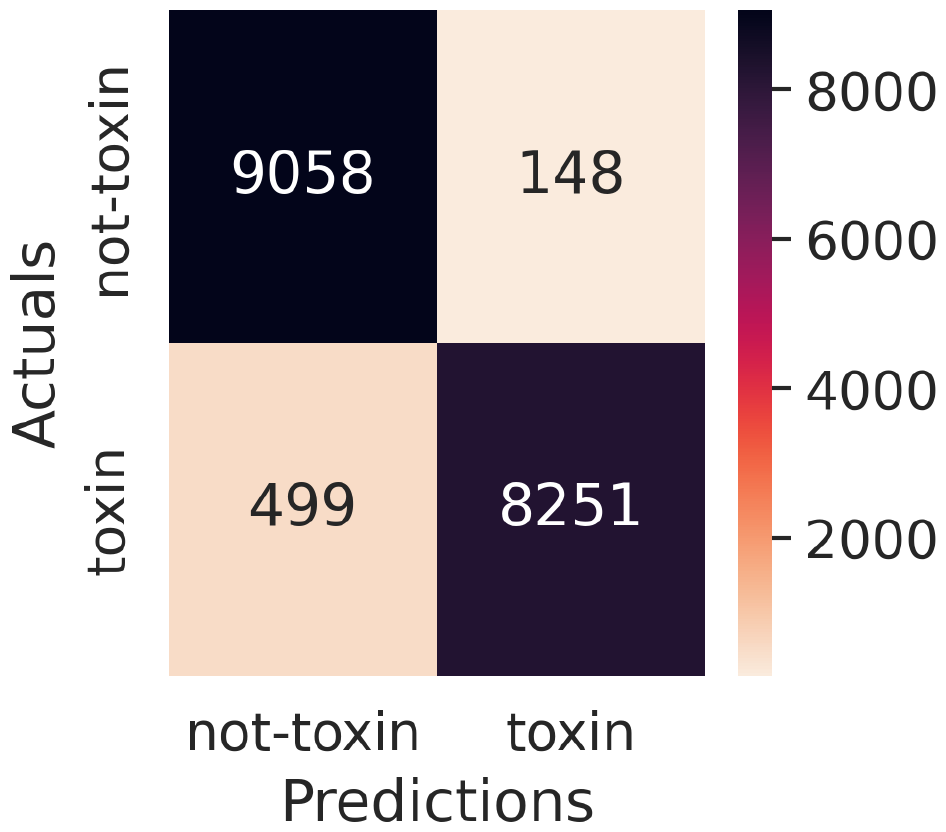
Confusion matrix for the BioLMTox prediction results on the validation dataset. All of the prediction results from Fig. 6 are aggregated (overlapping sequences between contemporary datasets are not duplicated) and enhanced with the additional curated data of the validation dataset.

### Embedding Analysis

Embedding analysis is an approach that can aid in model interpretability and dataset inspection. Embeddings refer to high dimensional features learned by the model during training. High level language aspects are represented by these high dimensional vectors that occupy a ‘semantic space’. Sequences that are more similar in meaning or function should have vectors that are closer together in this semantic space. BioLMTox embeddings are extracted from the hidden states of the last layer before the classification layers. As input passes through the model, more dimensions are added to the hidden states and the model should extract more abstract, complex and high level language features. Pooling techniques over these hidden states results in representations of the entire protein. The distances and clusters between these protein level representations may reflect differences and similarities in function, taxonomy or even structure.

Embeddings for the validation dataset were extracted with mean pooling (average of the hidden states over the token dimension) from the last hidden states of the fine-tuned BioLMTox and the pre-trained ESM-2 model using SentenceTransformers (45), producing vectors of size 1280 (3) for each sequence. To visualize these representations, their potential clustering, and the effect of the fine-tuning, Principal Component Analysis (PCA) was used to project both the ESM-2 pre-trained embeddings and BioLMTox embeddings into two dimensions. Emphasized in Fig.8, is the improved separation between the toxin and not-toxin clusters demonstrating the effectiveness of our fine-tuning methods and BioLMTox’s capability in extracting discriminative toxin features from protein sequences only.

While the embedding figures show BioLMTox has learned toxin features, the model’s predictive capabilities on the ToxinPred2 dataset (Table 2, Fig. 6) are not as performant. To investigate further, sequences in the validation dataset originating from ToxinPred2 are emphasized in the PCA projection of the BioLMTox embeddings (Fig. 9).

Figure 9 clearly shows that many of the highlighted Toxin-Pred2 toxin labeled datapoints (red) do not appear to be separated from the predominantly not-toxin cluster (blue). Additional datapoints are still only partially separated, ambiguous to either cluster. While the other datapoints do have some mingling with opposing clusters, the occurrence of this mingling is much less.

### BioLMTox: A General Toxin Classifier

Below we evaluate BioLMTox on the SOTA General Toxin Classifier Criteria:

### SOTA Dataset Criteria

#### 1. Domains of Life

Proteins in BioLMTox’s dataset consist of not just animal proteins but proteins of bacterial origin as well. Furthermore, stratified sampling was employed in dataset generation to make sure sequences of different taxonomic origin were represented. BioLMTox achieved high performance on datasets with these attributes, exhibiting capabilities as a general toxin classifier across different domains of life.

#### 2. Sequence Length

BioLMTox’s validation dataset consists of short peptides and longer proteins which allow classifier performance evaluation over the entire large range of sequence lengths. BioLMTox bias was not measured on specific sequence lengths.

#### 3. Sequence Similarity and Homologs

The capabilities of BioLMTox with regards to sequence similarity and homologs were not specifically evaluated in this study and remain an open opportunity for further investigation.

BioLMTox’s training dataset is much larger than many contemporary methods and so may require more resources per epoch than other contemporary methods. Furthermore, BioLMTox’s datasets include sequences from the UniProt TrEMBL database, which may potentially introduce noise into the datasets as they are unreviewed.

### SOTA Architecture Criteria

#### 4. Parameter and Feature Selection

BioLMTox is a single sequence model, which means additional features other than the sequence are not generated or needed for input. This is advantageous over alignment generated features that may not work on dissimilar sequences and often take time to generate. Furthermore, the embedding analysis demonstrates that BioLMTox is able to learn and extract discriminative toxin features on its own from just the protein sequence, removing the need for manual feature selection and threshold tuning. However, including additional features, while it may add additional complexity and overhead, can potentially increase the model’s performance.

#### 5. Computational efficiency and Overhead

BioLMTox leverages fine-tuning of a pre-trained model which should decrease the time and data iterations needed for model convergence compared to training the same architecture from scratch. However, as a transformer based model, BioLMTox does contain a significant number of parameters which increases the model’s complexity and memory footprint. The powerful capabilities of the fine-tuning methods described here, allow the model to be further fine-tuned on new data in the future and potentially reduce cumulative training time (time spent re-training a model and re-doing feature analysis/selection every time new data is incorporated into the training corpus). This means that novel and newly identified toxin sequences can be added to the model’s training corpus keeping the model contemporary with minimal additional intervention. Furthermore, BioLMTox is able to make sub-second classifications for single sequences, predicting labels for 16 sequences in 6.03 seconds, with no performance optimization, on a 12GB Nvidia RTX 3060ti.

#### Future Directions and Further Analysis

BioLMTox is a single sequence toxin classifier that achieves competitive results on eukaryotic and bacterial sequences of varying lengths. However, against the validation selection of the ToxinPred2 dataset, BioLMTox achieved less satisfactory results. The BioLMTox embeddings of the ToxinPred2 dataset (Fig. 9) and the conflicting labels uncovered during BioLMTox dataset creation, suggests labeling discrepancies between contemporary datasets. Metadata labels on sequence domain of life (Virus, Bacteria, Archaea etc), origin or function (neurotoxin, defensin, etc) were not provided in the ToxinPred2 main dataset downloaded files but remain a candidate for future identification and evaluation.

Further analysis and evaluation by grouping the ToxinPred2 dataset using taxonomy or GO annotation meta-labels might characterize these results. Using these same metadata labels, clustering in the embedding space could be further analyzed. In particular, the two subclusters for the toxin sequences in the embedding projections illustrated in Fig. 8 may be explained. Since one of these subclusters is predominantly made of sequences originating from BioLMTox dataset creation, we speculate that this cluster may be sequences from the bacterial domain of life or is a result of unreviewed sequences from the TrEMBL database, but without using meta-data labels we cannot confirm this speculation.

**Fig. 8.**
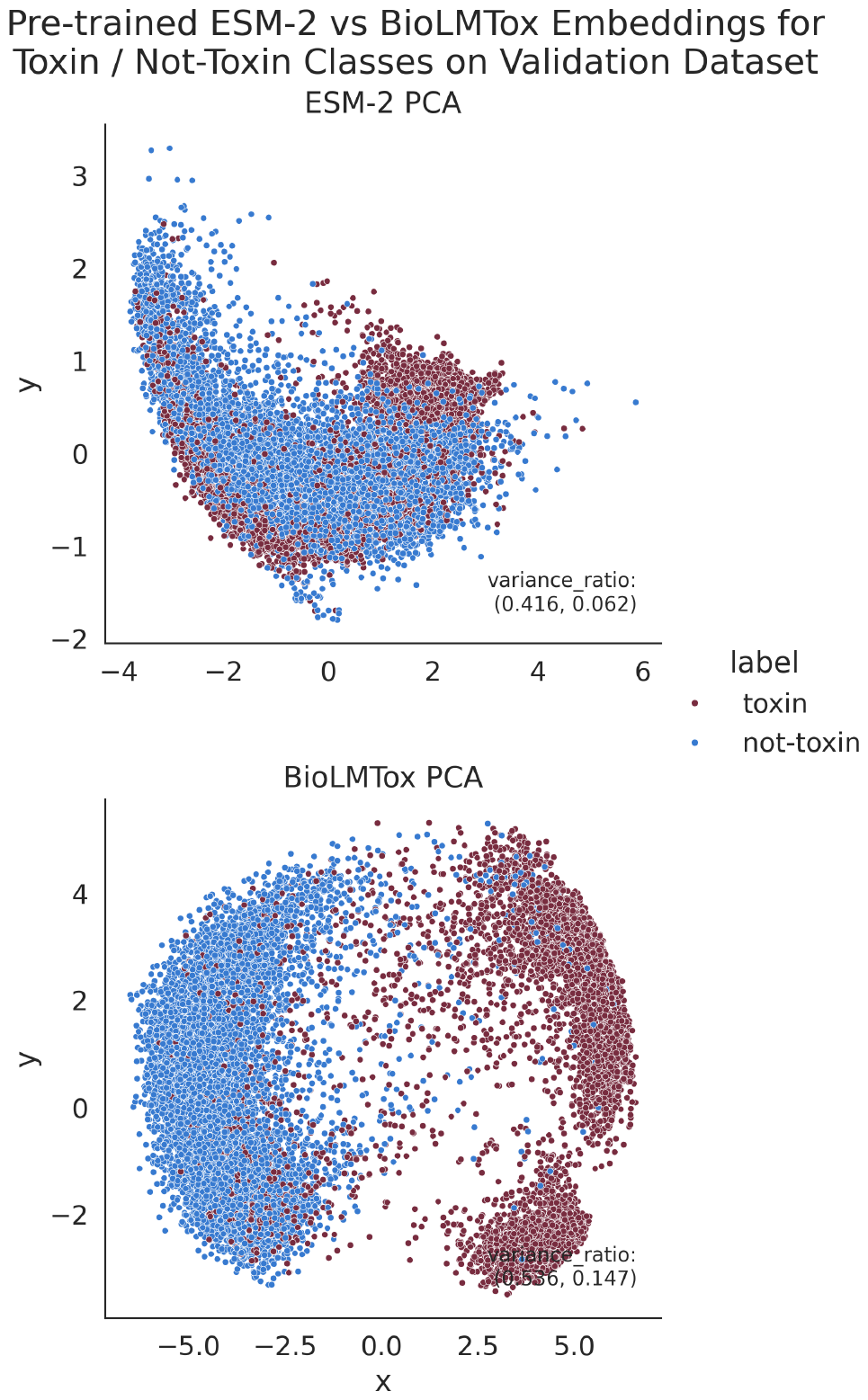
PCA projections of pre-trained ESM-2 and fine-tuned BioLMTox embeddings of BioLMTox’s validation dataset. Each point represents a protein sequence as represented by the models. Separation of the two classes is more evident for the BioLMTox embeddings demonstrating the fine-tuning’s effectiveness. The pretrained embeddings visually do not have clear separation while the BioLMTox embeddings do. Furthermore the variance ratio increases for both PCA components when projecting the BioLMTox embeddings. Within the projection of the BioLMTox embeddings, there also appear to be two separate subclusters for the toxin labeled sequences. One of these subclusters consists almost entirely of sequences unique to the BioLMTox dataset that do not exist in contemporary method datasets as emphasized in Fig. B.3. This suggests that this cluster may be related to the multiple domains of life and more explicit coverage of different taxonomies in the BioLMTox dataset.

**Fig. 9.**
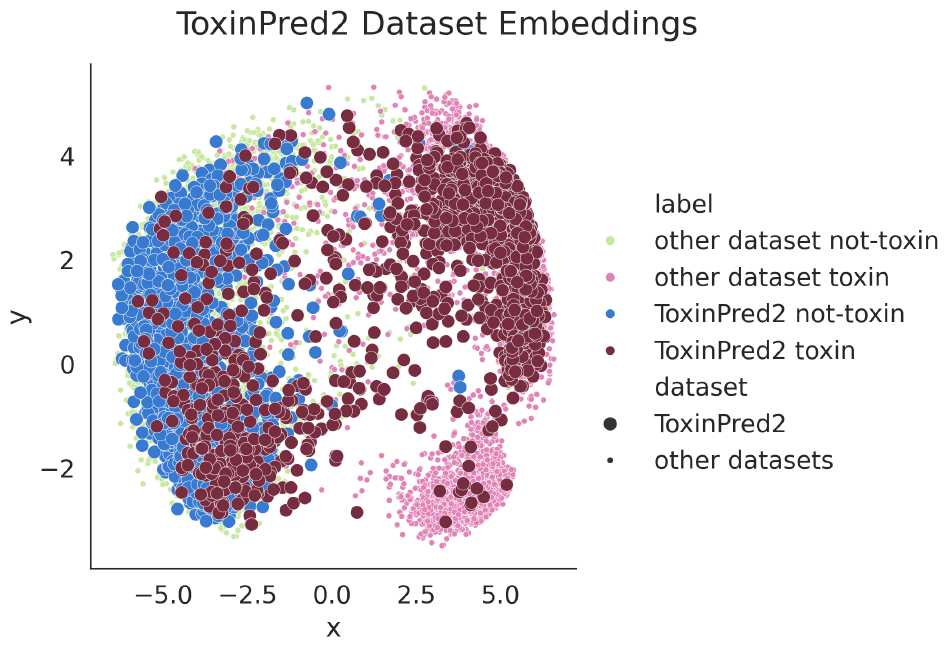
PCA projection of the BioLMTox embeddings for the validation dataset with the data from ToxinPred2 emphasized.

Conflicting labels also introduce another consideration for toxin classification. What defines a toxin and how should data be labeled to ensure quality training materials for machine learning models? With respect to biosecurity, toxins that are harmful to humans are the most significant concern. In the field of synthetic biology, protein sequences with antibiotic properties may be considered toxins and reduce yield if they are expressed in bacteria. Furthermore, prey specific toxins that may not be harmful to humans may still be of interest for academic research and therapeutic design. Additionally, there is a gradation to toxin effects that can be defined by lethal dose. This gradation may be another interesting component of toxin classification to explore.

Beyond just binary toxin classification there is also a target for multi-label toxin classification. Toxins can belong to subcategories such as neurotoxins, myotoxins and cardiotoxins. Training with finer grain predictions using these labels may produce interesting effects on embedding representations and lead to more informative predictions regarding toxin mechanism.

Furthermore, applying results in the developing field of Explainable AI (XAI) and Machine Learning (XML) (46, 47), to BioLMTox may lead to more biologically interpretable results, specifically in computationally determining mechanisms of protein toxicity. These methods may improve toxin domain and motif characterization and allow comparing model explanations that can be validated with experimental observations.

The SOTA general toxin classifier criteria proposed here are heuristic and no quantitative thresholds for what methods should be considered SOTA were explicitly proposed. This was partly due to the immediate need to first define what capabilities a ‘SOTA’ method should display with respect to increasing dual-use concerns, and partly in order to directly compare BioLMTox against contemporary methods on their specific datasets. Considering BioLMTox’s high performance and straightforward application as an initial attempt towards these criteria, future studies with BioLMTox using more explicit datasets that include carefully selected out of distribution (OOD) datasets and blind test sets to determine quantitative benchmarks for each of these specific criteria are compelling. Furthermore, since BioLMTox demonstrates that even a simple fine-tuning architecture using a pre-trained language model is effective for toxin classification, a relevant and important biological problem, there is motivation to test more complex or nuanced architectures and fine-tuning techniques.

### Benchmark Dataset

For the purpose of direct comparison with existing toxin classifier methods and to satisfy several of the SOTA general toxin classifier criteria, we propose the BioLMTox dataset as a benchmark toxin classification dataset. The BioLMTox validation dataset contains data from five contemporary datasets: ToxinPred2, CSM-Toxin, ToxDL, ATSE and TOXIFY, which enables comparison with nine methods. The CSM-Toxin dataset contains sequences up to 15639 amino acids (AAs) in length, the ATSE dataset contains peptide sequences between 6 and 50 AAs and TOXIFY contains sequences up to 500 AAs. These datasets provide ample validation data at different length distributions. Furthermore, the data from UniProt and UniRef50 selected during BioLMTox dataset creation and the data from Toxin-Pred2 consist of protein sequences from more than just eukaryotic organisms allowing for validation on multiple domains of life. Furthermore, we curate the data removing ambiguously labeled sequences between contemporary datasets, and discuss potential discrepancies for toxin definitions between the contemporary datasets.

This unified dataset will be available shortly and can be used to evaluate and facilitate future general toxin classification efforts.

## Conclusions

In this study, we reviewed applications of *in-silico* toxin classification, reviewed current contemporary methods for toxin classification, identified five heuristic criteria for SOTA general toxin classification, discussed contemporary methods’ strengths and weaknesses with respect to these criteria, compiled a unified dataset for toxin classification that satisfies several of the proposed SOTA dataset criteria, and proposed BioLMTox, a single sequence toxin classifier trained on eukaryotic and bacterial protein and peptide sequences. BioLMTox demonstrates capacity as a general toxin classifier: it does not use additional features other than the sequence, has a simple linear classification head, is based on a medium size ESM-2 pre-trained model, and performance exceeds or is comparable to contemporary methods on sequences ranging from 5 to 15639 amino acids in length. BioLMTox exemplifies the proficiency of simple fine-tuning for specific computational biology problems and has potential applications in synthetic biology, as a biosecurity screen and for protein, peptide and drug research.

## BioLMTox Availability

BioLMTox will be available and hosted at [BioLM](https://biolm.ai/) shortly.

## Acknowledgments and Contributions

We are grateful to Ahmad Qamar for his insightful suggestions on structuring the manuscript.

ACKNOWLEDGEMENTS

## Appendix A Datasets

All UniProt sequences used were downloaded in tsv file format. The UniRef50 database was downloaded in fasta file format. All UniRef50 sequences used were labeled using the Tax ID extracted from the fasta file and then the corresponding lineage name and rank from the [UniProt Taxonomy Database](https://www.uniprot.org/taxonomy?query=%2A). All sequences obtained from UniProt and UniRef50 databases were processed to remove duplicate sequences, sequences with non standard amino acid codes ‘B’, ‘J’, ‘O’, ‘U’, ‘X’, ‘Z’, and sequences with ‘virus’, ‘Virus’, ‘archaea’ or ‘Archaea’ in their taxonomic lineage or superkingdom (domain of life). Other contemporary model datasets were downloaded via github, another repository or web-app, if available, and aggregated into a combined dataset.

### A.1. Contemporary Datasets

For comparison to contemporary toxin classification methods, datasets were extracted from the pre-existing dataset splits of the models listed in Table 1. Several of these methods, such as ToxVec and TOXIFY, share datasets. The datasets used were from [TOXIFY, ToxDL, ATSE, ToxinPred2](https://github.com/xypan1232/ToxDL/tree/master/datasets) and CSM-Toxin.

All of the training datasets for these methods were combined into one training dataset split: C-T (Contemporary Training), and likewise the validation, test or blind test datasets were combined into one validation dataset split: C-V (Contemporary Validation) (Fig.1).

Any of the sequences in C-T that were in C-V were removed from training. All of the sequences from SO-T that were greater than 2048 in length (twice BioLMTox’s context window) were discarded. The CSM-Toxin and TOXIFY methods’ training datasets had large imbalances of not-toxin labeled sequences that were downsampled to equal the number of toxin labeled sequences in each of their respective datasets prior to combining with the other training datasets when creating C-T. Note: the explicit splits for the ToxinPred2 training and validation were not provided so a 8:2 random split of ToxinPred2’s main dataset was used to create training and validation datasets as detailed in the original study (23). The ToxDL validation dataset was also included, along with the ToxDL’s primary test split, in the C-V.

There were several sequences with contradicting labels between datasets. These ambiguously labeled sequences were removed from both the C-T and C-V datasets. Ambiguous labels would have made the breakdown of BioLMTox’s evaluation on these validation datasets unreasonable, as correct predictions in one dataset would be incorrect predictions for the conflicting dataset. Ambiguously labeled datapoints are visualized in Fig. B.3. Most of these conflicting labels occurred between ToxinPred2 and other datasets. It is worth noting that these datasets were created at different times, and some proteins may have been recently identified as toxins. Furthermore, the definitions of what constitutes a toxin may have differed when creating these datasets; For example, datasets that only selected animal venoms as their defined toxins as opposed to datasets that also included bacterial toxins in their definition of toxins. The definition of a toxin can influence whether sequences that are toxic to plants or bacteria are included with proteins that are toxic to animals or whether UniProt keywords, GO annotations or a combination of both are used when labeling data. One of the ambiguously labeled sequences, [M-ectatotoxin-Eb2c](https://www.uniprot.org/uniprotkb/C0HK47/entry) is labeled with the antibiotic UniProt keyword but is missing the toxin keyword. However, the GO annotations for this entry have the toxin activity annotation. ATSE labeled this sequence as a not-toxin while TOXIFY labeled it as a toxin.

### A.2. UniProt and UniRef Sequences

A toxin dataset was created by downloading toxin sequences from UniProt labeled with “KW 0800: Toxin” and the addition of “NOT virus” and “NOT archaea” with this specific query. Then sequences also in C-V and sequences greater than 1024 or less than 6 AA’s in length were discarded. A not-toxin dataset was created by discarding all sequences greater than 1024 and less than 6 AA’s from the UniRef50 dataset and then further removing all of the toxins sequences downloaded from UniProt and the sequences in C-V. Then a stratified sampling using taxonomic class as the strata was performed on this not-toxin dataset to reduce the size of the dataset to a manageable size (a similar size as the toxin dataset) for training and to ensure sequences from as many taxonomic classes as possible were included in the reduced dataset.

### A.3. Validation Dataset

An equal number of sequences to those in C-V (A.1) were randomly selected from the toxin and not-toxin datasets created in (A.2) and added to C-V to to form the base for BioLMTox validation dataset. Then an additional 2500 toxin labeled sequences from the toxin dataset detailed in Section A.2 were added. This balanced the toxin and not-toxin classes better enabling accuracy to be a hyperparameter tuning objective. Any duplicate sequences were discarded.

### A.4. Comparison Training Dataset

Non redundant sequences from C-T (A.1) were added to the remaining toxin and not-toxin sequences (those were not selected for the validation dataset (A.3) to create the training dataset.

As shown in Fig. 2, The majority of data in BioLMTox datasets was selected from UniProt and UniRef as compared to the data selected from contemporary methods.

## Appendix B Additional Figures

**Fig. B.1.**
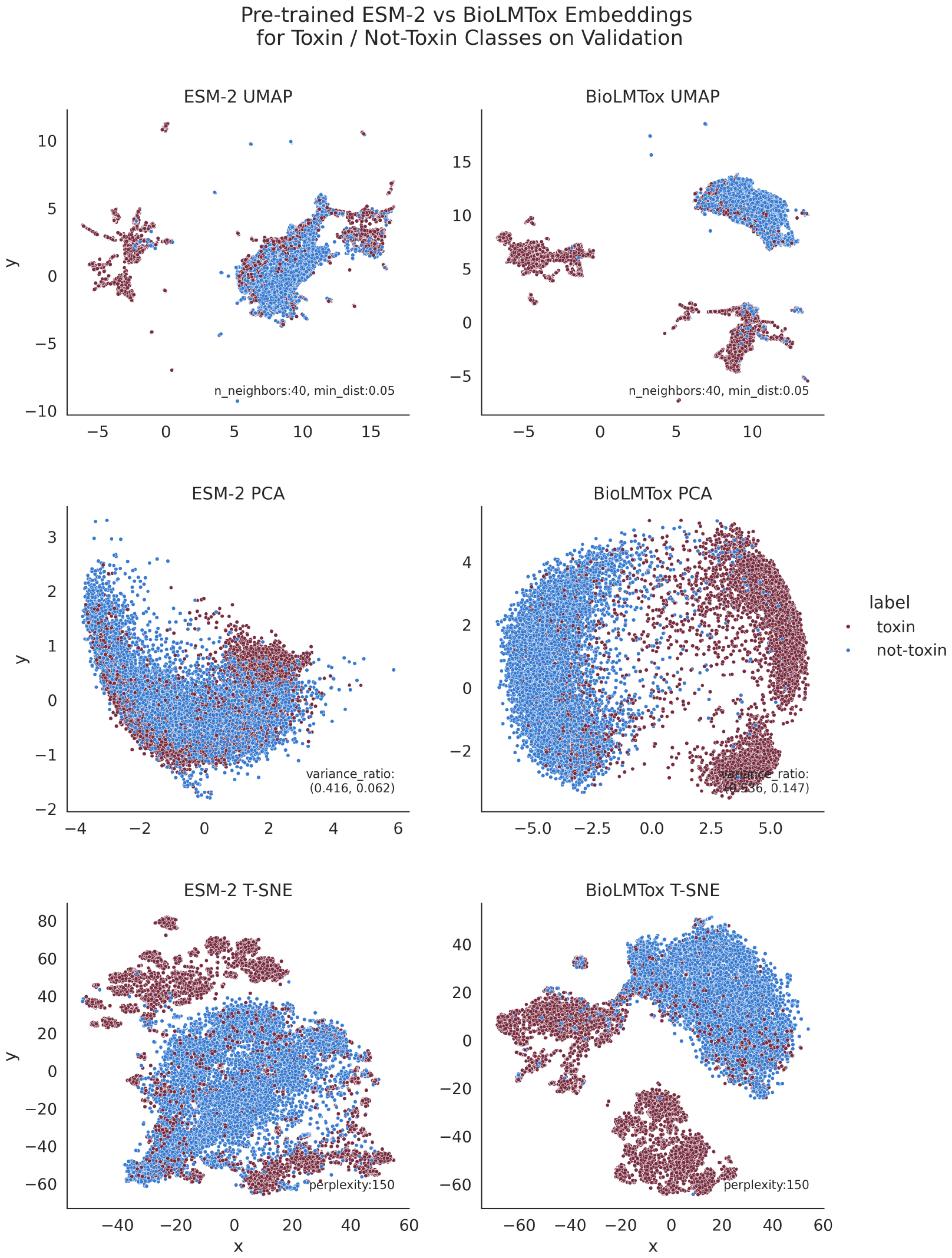
Different projection methods including Uniform Manifold Approximation and Projection (UMAP), Principal Component Analysis (PCA) and t-distributed Stochastic Neighbor Embedding (T-SNE) visualizing pre-trained ESM-2 and fine-tuned BioLMTox embeddings for the validation. All projection methods for the BioLMTox embeddings show what appear to be two separate subclusters for the toxin labeled sequences. Figure B.3 shows that not many sequences originating from other methods are contained in this cluster showing that these datapoints originated with BioLMTox dataset construction.

**Fig. B.2.**
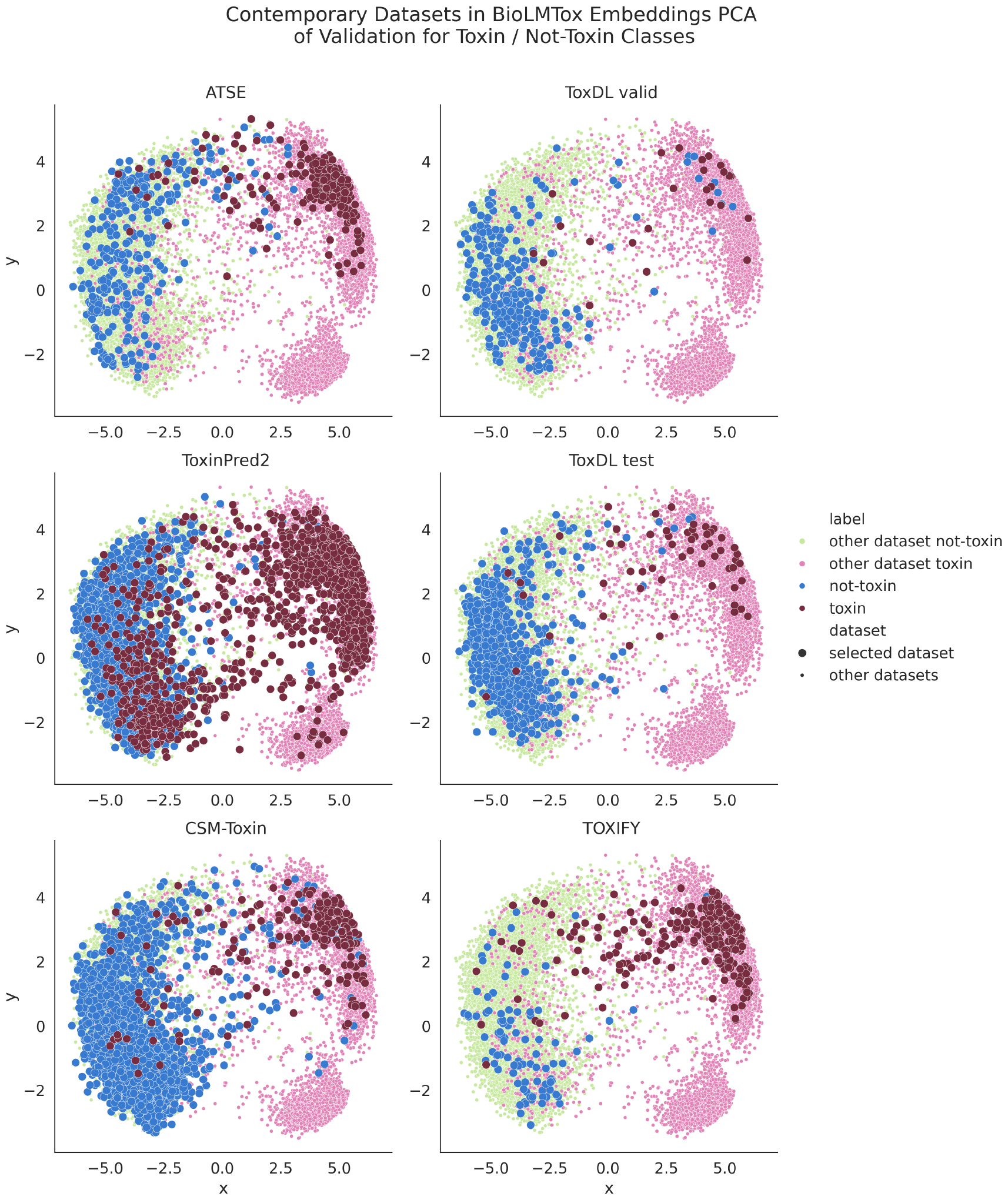
PCA projection of the BioLMTox embeddings for the validation dataset with the data from each comparison method emphasized. Each subplot has larger blue not-toxin and red toxin colored datapoints that correspond to data from the different contemporary methods. The smaller green not-toxin and pink toxin datapoints correspond to data in the validation that does not belong to the selected highlighted method. Most of the highlighted points maintain good separation into the opposing clusters with the exception of ToxinPred2 which has significant mingling of toxins into the predominantly not-toxin visual cluster. It should be noted that the greater frequency of unseparated toxins for the ToxinPred2 dataset compared to the other contemporary datasets may be in part because it is a larger dataset. One of the apparent toxin subclusters does not contain many sequences from contemporary datasets (only toxins from ToxinPred2) indicating that these datapoints originated with BioLMTox dataset construction and suggesting that these sub clusters may relate to different domains of life (one significant difference between ToxinPred2 and BioLMTox datasets compared with other contemporary methods).

**Fig. B.3.**
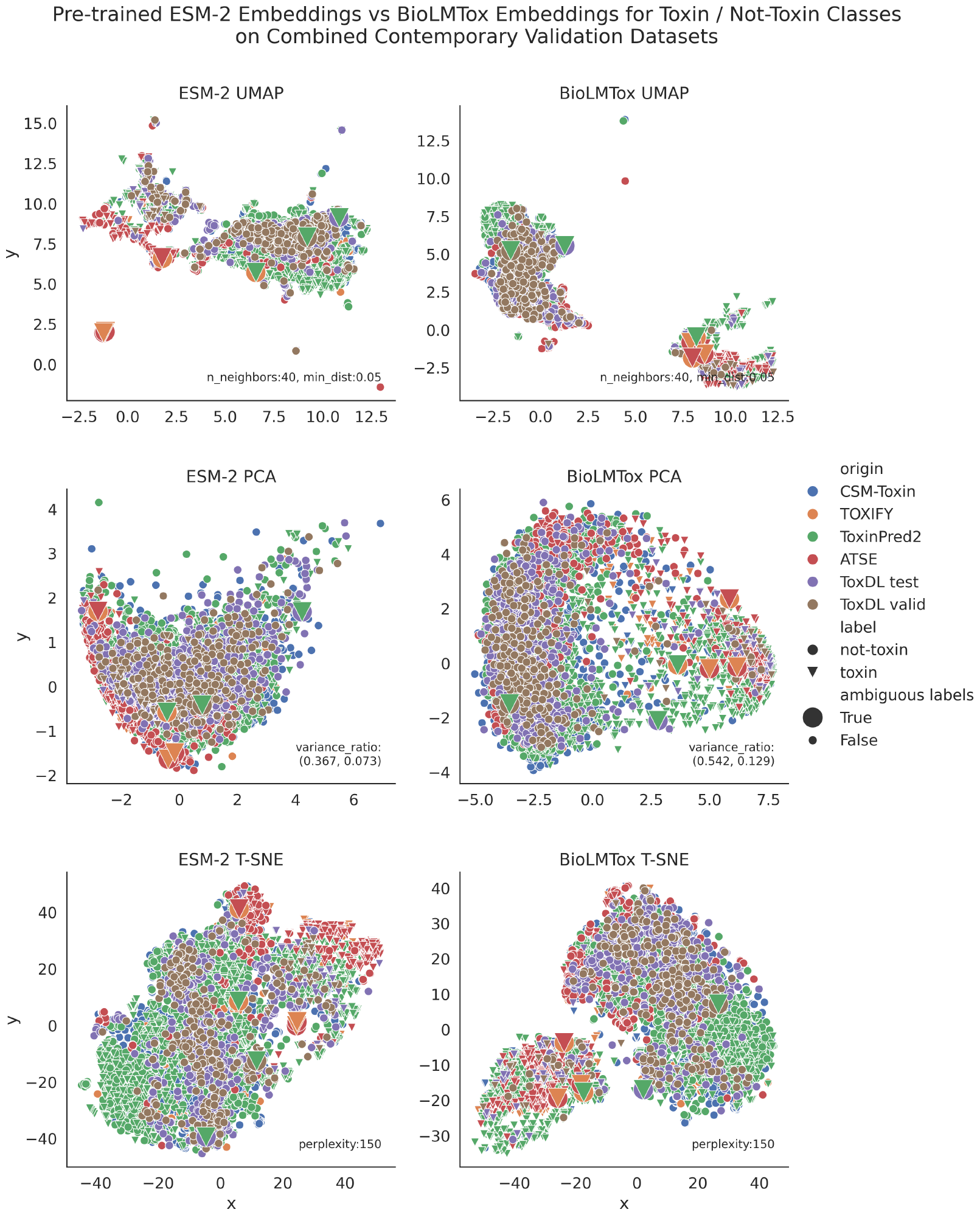
The pre-trained and BioLMTox embeddings for the BioLMTox validation dataset consisting of only sequences from contemporary methods are visualized along with the ambiguously labeled datapoints discovered during BioLMTox dataset creation. There are 6 pairs of these ambiguously labeled datapoints emphasized with larger point markers. Additionally, the contemporary sequences still exhibit a similar separation between toxin and not-toxin classes as the full validation dataset when comparing the pre-trained and BioLMTox embedding space as demonstrated by the increase of explained variance of the first two principal components which approaches the values shown in Fig. B.1.

